# Rapid evolution drives the rise and fall of carbapenem resistance during an acute *Pseudomonas aeruginosa* infection

**DOI:** 10.1101/2020.08.10.243741

**Authors:** Rachel Wheatley, Julio Diaz Caballero, Natalia Kapel, Angus Quinn, Ester del Barrio-Tofiño, Carla López-Causapé, Jessica Hedge, Gabriel Torrens, Thomas Van der Schalk, Basil Britto Xavier, Felipe Fernández-Cuenca, Angel Arenzana, Claudia Recanatini, Leen Timbermont, Frangiscos Sifakis, Alexey Ruzin, Omar Ali, Christine Lammens, Herman Goossens, Jan Kluytmans, Samir Kumar-Singh, Antonio Oliver, Surbhi Malhotra-Kumar, Craig MacLean

**Author notes:** these authors contributed equally to the presented work (Rachel Wheatley, Julio Diaz Caballero).

## Abstract

It is well established that antibiotic treatment selects for resistance in pathogenic bacteria. However, the evolutionary responses of pathogen populations to antibiotic treatment during infections remain poorly resolved, especially in acute infections. Here we map the evolutionary responses to treatment in high definition through genomic and phenotypic characterization of >100 isolates from a patient with *P. aeruginosa* pneumonia. Antibiotic therapy (meropenem, colistin) caused a rapid crash of the *P. aeruginosa* population in the lung, but this decline was followed by the spread of meropenem resistance mutations that restrict antibiotic uptake (*oprD*) or modify LPS biosynthesis (*wbpM*). Low fitness strains with high-level meropenem resistance (*oprD*) were then replaced by high fitness strains with ‘anti-resistance’ mutations in the MexAB-OprM efflux pump, causing a rapid decline in resistance to both meropenem and a collateral loss of resistance to a broad spectrum of antibiotics. In contrast, we did not observe any evolutionary responses to antibiotic treatment in the intestinal population of *P. aeruginosa*. Carbapenem antibiotics are key to the treatment of infections caused by Gram negative pathogens, and our work highlights the ability of natural selection to drive both the rapid rise and fall of carbapenem resistance during acute infections.

## Introduction

Antibiotic resistance has emerged as a serious threat to public health by increasing the mortality rate and economic costs associated with bacterial infections [1]. Treating patients with antibiotics selects for resistant strains [2, 3], and the emergence of resistance during treatment is associated with poorer outcomes in terms of patient health [1, 4]. Following treatment, resistance in patients typically returns to baseline levels, although there is considerable heterogeneity in the rate of decline for different microbe/antibiotic combinations [5, 6]. Although this link between antibiotic treatment and resistance is straightforward and intuitive, the dynamics of the rise and fall of antibiotic resistance during infections remain poorly understood. Progress in this area has largely come from studies that have used longitudinal sampling of patients to study the emergence of resistance during long-term chronic infections associated with diseases such as cystic fibrosis and tuberculosis [7-13]. Although these studies have produced important insights into resistance, short-term acute infections, such as hospital-acquired infections by opportunistic and commensal pathogens, represent a very important burden of AMR [14].

Here we investigate the evolutionary responses of bacteria to antibiotic treatment through intensive sampling of a single mechanically ventilated patient before, during, and after treatment for a hospital-acquired *Pseudomonas aeruginosa* pneumonia. *P. aeruginosa* is an opportunistic pathogen that is a relatively common cause of nosocomial infections, particularly in immunocompromised patients [15-18], and *Pseudomonas* pneumonia infections of the lung are associated with a very high mortality rate (i.e. 30% for ventilator-associated pneumonia) [17]. *P. aeruginosa* infections are difficult to treat because this pathogen has high levels of intrinsic antimicrobial resistance due to low outer membrane permeability combined with a large repertoire of both intrinsic resistance mechanisms, such as MexAB-OprM multidrug efflux pump, and acquired resistance mechanisms, including chromosomal mutations and acquisition of mobile resistance genes [19-21].

To understand the responses to antibiotic treatment, we intensively sampled *P. aeruginosa* from the lung (endotracheal aspirate) and gut (peri-anal swabs) at regular intervals over three weeks. We collected 12 isolates from each sample, and we used whole-genome sequencing and phenotype assays (resistance profiling, fitness) on over 100 isolates to understand the population-level responses to antibiotic therapy. By combining this population biology data with clinical data, we have been able to produce a high-resolution characterization of bacterial responses to antibiotic treatment.

## Results and Discussion

### Clinical data

A 60-year-old female patient was admitted to the intensive care unit (ICU) of the Virgen Macarena tertiary care hospital in Seville, Spain with a primary diagnosis of haemorrhagic shock. The patient was intubated and started on mechanical ventilation, and given prophylactic treatment with amoxicillin/acid clavulanic (1000 mg/200 mg IV q8h). After 72 hours of ICU admission, informed consent was obtained and the patient was enrolled in the ASPIRE-ICU study (day 1) [22]. On day 1, the titre of *P. aeruginosa* in the endotracheal aspirate (ETA) was high at 10^6^ colony-forming units per ml (CFU/mL) and *P. aeruginosa* were the only culturable bacteria that were detected in ETA samples (Figure 1). A clinical diagnosis of pneumonia was established by the treating physician on day 2 and the patient was treated with piperacillin/tazobactam (4 g/0.5 g IV q8h for 2 days), meropenem (1 g IV q8h for 2 days) and colistin (3 million IU IV q8h for 13 days) (Figure 1).

**Figure 1.**
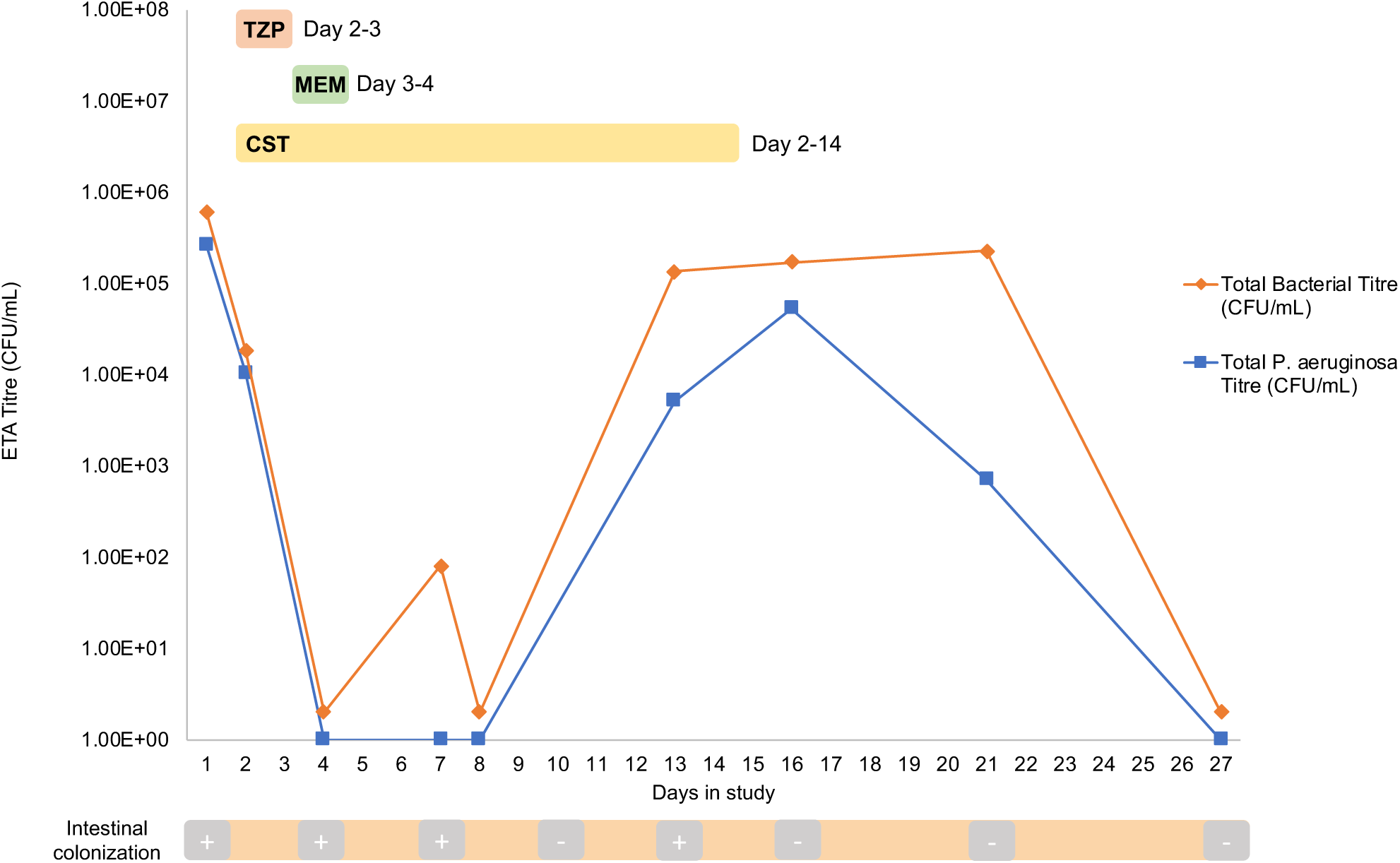
Clinical timeline of patient. The titre of *P. aeruginosa* in the lung was assessed by plating out samples of endotracheal aspirate (ETA) on *Pseudomonas* selective agar and total bacterial titre was determined by plating out ETA on blood agar. Intestinal colonization was assessed semi-quantitatively by plating out samples from peri-anal swabs on *Pseudomonas* selective agar; (+) indicates *Pseudomonas* detection at this sampling point, (-) indicates no *Pseudomonas* detection at this sampling point. Piperacillin/tazobactam (TZP), meropenem (MEM) and colistin (CST) were delivered intravenously at the following doses: TZP; 4 g/0.5 g IV q8h for 2 days, MEM; 1 g IV q8h for 2 days, CST; 3 million IU IV q8h for 13 days. Day 1 in study is 72h after ICU admission and corresponds to the first day of patient informed consent.

The microbiological response to antibiotic treatment was dramatic: the titre of *P. aeruginosa* in the lung declined by >10 fold between day 1 and 2, and fell to below the assay detection limit of 40 CFU/mL by day 4. The decline in *Pseudomonas* titre was associated with improved patient health: between day 2 and 7 the sequential organ failure assessment score (SOFA) declined from 14 to 9, and the Clinical Pulmonary Infection Score (CIPS) declined from 8 to 4. In spite of the initial success of antibiotic treatment, culturable bacteria were detected in ETA samples at day 8, and *P. aeruginosa* began to rebound, eventually achieving a density of 10^4^ -10^5^ CFU/mL. Nevertheless, no new episodes of clinical pneumonia were reported and no new antibiotics were administered. Ventilator support was withdrawn on day 23, no *Pseudomonas* was detected in ETA samples taken on day 27, and the patient was discharged from ICU on day 31 with a SOFA score of 1 and a CPIS of 0.

We found evidence of intestinal colonization by *P.* aeruginosa on admission to ICU, as measured by growth from peri-anal swabs. Qualitative measures of the intestinal abundance of *P. aeruginosa* remained essentially constant up until day 13, but no growth of *P. aeruginosa* was detected in peri-anal swabs that were taken from day 16 onwards (Figure 1). In total, we collected 107 isolates of *P. aeruginosa* from ETA samples (n=59) and peri-anal swabs (n=48).

### Resistance phenotyping

To understand the role of antibiotic treatment in the decline and subsequent expansion of the pulmonary population of *P. aeruginosa*, we measured the resistance of lung isolates to meropenem, piperacillin/tazobactam, and colistin (Figure 2). Isolates from early time points (day 1-2) had high levels of piperacillin/tazobactam resistance (MIC>256mg/L) that were well above the clinical breakpoint (16mg/L), suggesting that piperacillin/tazobactam treatment is unlikely to have had any effect on *P. aeruginosa* (Figure 2A). In contrast, levels of resistance to meropenem (mean MIC=9.61 mg/L; s.e.m=.677; n=24) were close to the published EUCAST clinical breakpoint concentration (8 mg/L), and colistin MICs (mean=0.5 mg/L; s.d=0; n=24) were below the clinical breakpoints (2mg/L), suggesting that these two antibiotics contributed to the collapse of the pulmonary population of *P. aeruginosa*.

**Figure 2.**
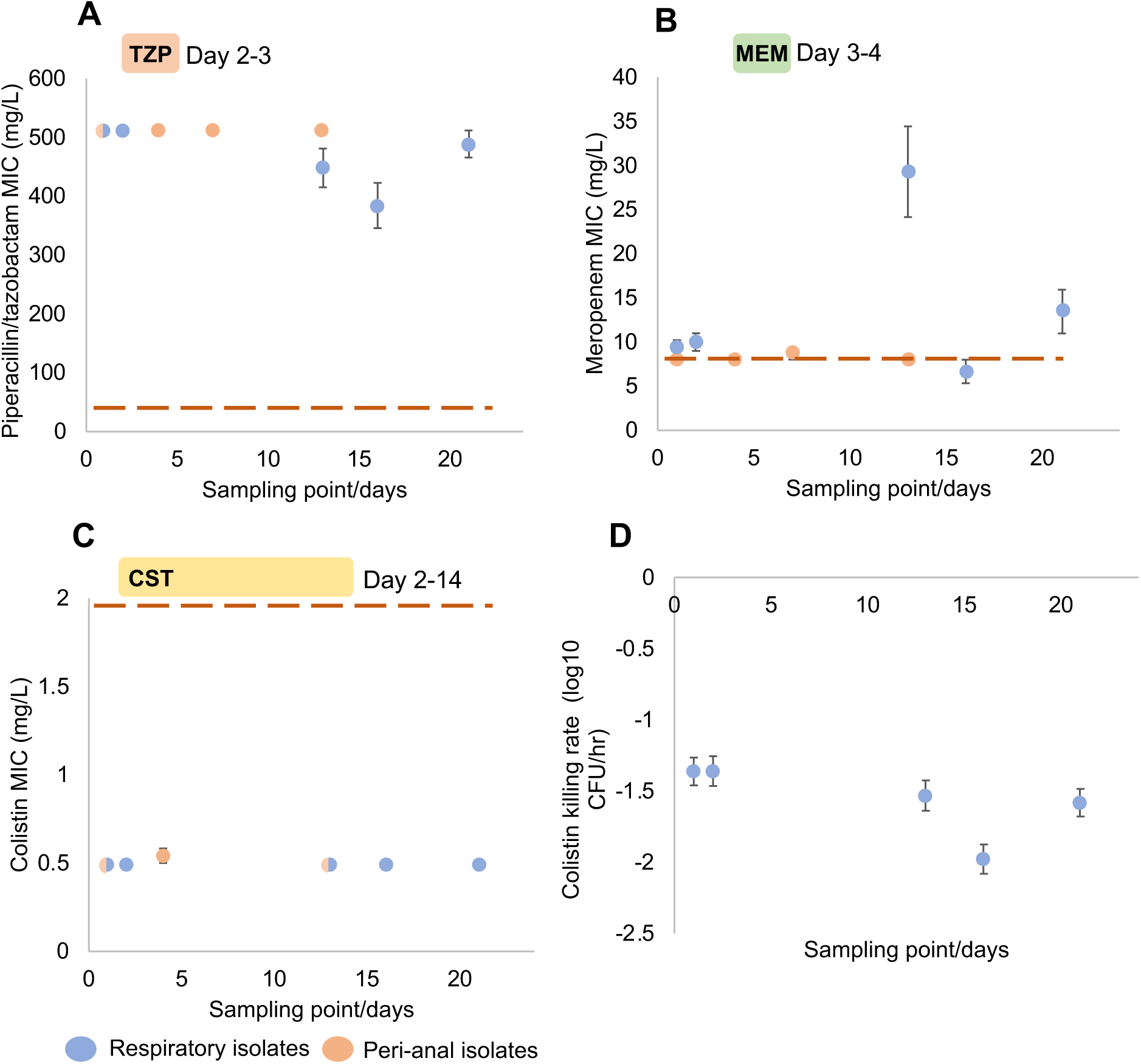
Mean +/- S.E.M resistance (MIC measured via broth microdilution) to A) piperacillin/tazobactam (TZP), B) meropenem (MEM) and c) colistin (CST) for both the respiratory and peri-anal isolates collected over 3 weeks in ICU. The red dashed line represents the EUCAST clinical breakpoint (01/01/2019 edition); 16 mg/L (TZP), 8 mg/L (MEM) and 2 mg/L (CST). D) The colistin killing rate (mean +/- s.e.m) of cells following treatment with 2mg/L of colistin. Sampling points (/days) corresponds to the clinical timeline detailed above.

The pulmonary population of *P. aeruginosa* began to recover while the patient was still being treated with colistin, suggesting that colistin resistance may have increased during the infection. However, the MIC of isolates from late time points (days 13-21) did not increase (mean MIC=0.5 mg/L; s.e.m=0; n=35), demonstrating that genetically elevated levels of colistin resistance did not evolve during the infection (Figure 2C). To test for subtle changes in colistin resistance that cannot be captured by MIC assays, we measured the viable cell titre of cultures of pulmonary isolates following treatment with 2mg/L of colistin. The density of viable cells declined very rapidly following colistin treatment (mean=-1.57 log10 CFU.ml-1.hr-1; s.d=.57), and we found no evidence that colistin tolerance increased following antibiotic treatment (Figure 2D).

In contrast to colistin and piperacillin/tazobactam, we found dynamic changes in meropenem resistance over the course of the infection. Meropenem resistance increased following antibiotic treatment (day 13: mean MIC=29.33 mg/L; s.e.m=1.04; n=12), suggesting that treatment drove the rapid evolution of resistance. This response to meropenem treatment was short-lived, and levels of resistance rapidly declined to baseline levels at day 16 (mean MIC=6.66 mg/L; s.e.m=1.33; n=12) and day 21 (mean MIC=13.45 mg/L; s.e.m=2.48; n=11).

In contrast, semi-quantitative measures of the prevalence of *P. aeruginosa* in peri-anal swabs did not decline during antibiotic treatment, and all of the peri-anal isolates had the same antibiotic resistance profile, apart from the fact that a single isolate had a marginally higher resistance to colistin (MIC=1 mg/L).

### Sequencing

To better understand the rise and fall of resistance, we used a combination of short and long-read sequencing to comprehensively characterize the *P. aeruginosa* genetic diversity found in the patient. Initially, we used long-read sequences generated by PacBio to generate a high-quality reference genome for one of the day 1 pulmonary isolates (Figure 3A). This ST17 reference isolate has a large genome (7,008,585 bp) and a 40Kb plasmid, p110820, that carries genes conferring resistance to aminoglycoside [*aacA4*], ß-lactam [*bla-Oxa10*], and sulphonamide [*sul1*] antibiotics. The reference genome contains many mutations that are associated with antibiotic resistance, including target modification mutations that confer resistance to fluoroquinolones (*gyrA* T83I *and parC* S87L) and mutations in repressors of the AmpC ß -lactamase (*ampD* H98Y) and the MexAB-OprM multidrug efflux pump (*nalD*nt58Δ2) [20, 23]. Moreover, the overexpression of *ampC* (37.6 ± 21.7 fold) and *mexAB-OprM* (20.8± 17.4 fold) in four randomly chosen isolates relative to the PA01 reference strain was confirmed with Real Time RT-PCR. This clone of ST17 has previously been reported as the cause of a nosocomial outbreak at the Virgen Macarena hospital [24].

**Figure 3.**
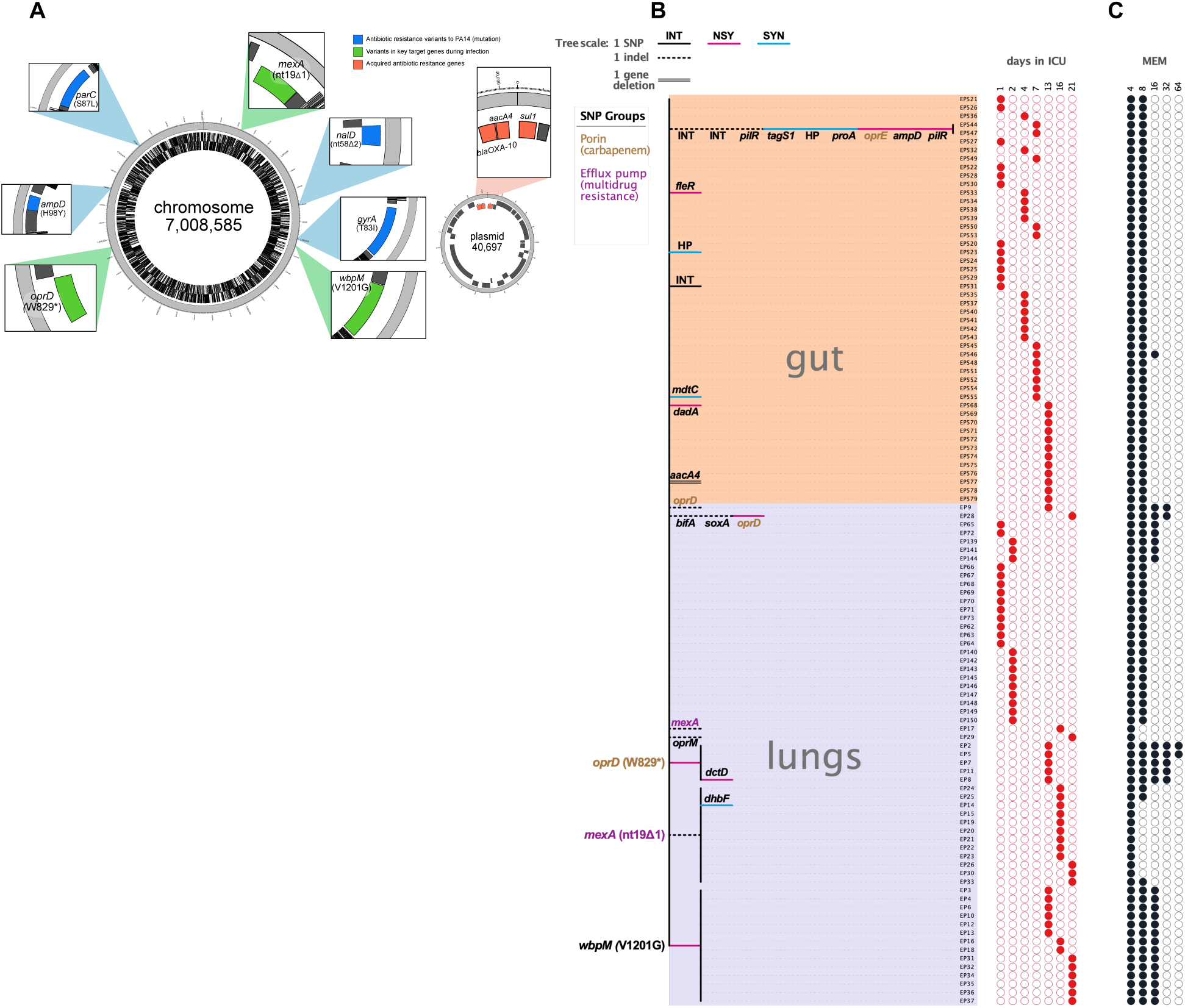
Assembly of a closed reference genome for the ST17 clone that initiated lung infection. A) Annotations show SNPs and acquired resistance genes on the plasmid and chromosome. SNPs are annotated relative to the PA14 reference genome. B) Neighbour-joining tree showing SNPs and indels in lung and gut isolates, highlighting three lineages that evolved during respiratory infection. C) Meropenem resistance score (MIC in mg/M).

To identify mutations that arose during infection, we mapped short reads from all 107 isolates to this closed reference genome (Figure 3B). Using this approach, we found a small number of SNPs (n=16) and indels (n=9), most of which occurred as singletons (n=13). To identify the variable genetic content among our isolates (the part of the genome found only in some isolates), we compared the genetic composition of each isolate against the gene content of all isolates, and validated the potential variable genome by mapping sequencing reads to the sequences of genes in these regions. The only evidence of changes in genome composition that we found was the loss of the plasmid carried *aacA4* aminoglycoside resistance gene in a single intestinal isolate.

### The rise of meropenem resistance

All the lung isolates from days 1 and 2 were clonal ST17, demonstrating that the pneumonia infection was the result of the recent expansion of a single dominant clone in the lung, hereafter called the ancestral strain. The recovery of the *P. aeruginosa* population following antibiotic treatment was associated with an increase in meropenem MIC at day 13. At a genetic level, this was driven by the replacement of the ancestral strain by mutants with elevated meropenem resistance.

Mutational resistance to meropenem in *P. aeruginosa* is often attributable to mutations in the outer membrane porin *oprD* [25]. We detected 3 independent *oprD* mutations, including a premature stop codon (W829*) and a frameshift (nt1206Δ5), and isolates with these mutations had elevated meropenem resistance (mean MIC=41.1 mg/L; s.e.m=5.9; n=7) compared to isolates of the ancestral strain (Dunnett’s test; P<.0001). In total, 5/12 of the day 13 isolates carried an *oprD* mutation, but mutations in this gene were not detected at later time points, except for a single isolate carrying an *oprD* frameshift mutation.

Almost all of the remaining (6/12) day 13 isolates carried a mutation in a gene (*wbpM* [V1201G]) that is part of a lipopolysaccharides biosynthesis operon [26] and has been previously implicated in resistance to ß-lactam antibiotics, including meropenem [27]. In line with these results, isolates with the *wbpM* mutation had 2-fold higher levels of resistance (mean MIC=16mg/L; s.e.m=0; n=14) than isolates of the ancestral strain (Dunnett’s test; P=.0045). To further validate this association, we tested a *wbpM* transposon mutant of *P. aeruginosa* PA14 for elevated meropenem resistance. The transposon mutant was associated with a 2x increase in meropenem resistance, confirming that the loss of this gene provides a marginal increase in meropenem resistance.

### The fall of meropenem resistance

Late in the infection (days 16-21), levels of meropenem resistance declined back to baseline levels. At a genetic level, this was driven by the replacement of *oprD* W277 mutants by isolates with mutations in the MexAB-OprM efflux pump. MexAB-OprM is a broad-spectrum antibiotic efflux pump that is often upregulated in *P. aeruginosa* clinical isolates due to mutations in the *nalD* repressor [20, 23]. Strikingly, we observed 3 independent losses of this efflux pump via frameshift mutations in both *mexA* and *oprM*. As expected, isolates with MexAB-OprM mutations had reduced resistance to meropenem (mean MIC=4.85mg/L; s.e.m=0.45; n=14) relative to isolates of the ancestral strain (Dunnett’s test; P=.040). MexAB-OprM mutants were present at a high frequency at both day 16 (10/12) and 21 (5/11). The decline in meropenem resistance observed after day 13 (Figure 2B) was predominantly driven by the replacement of the dominant *oprD* W277 mutant by the *mexA* nt19Δ1 mutant.

### Gut isolates

In contrast to the rapid evolution observed in the lung, we found no evidence for evolutionary adaptation in gut isolates. Five isolates carried singleton SNPs, and the only SNP in a gene associated with an annotated role in antibiotic resistance (*mdtC*) did not lead to a change in the amino sequence, explaining why we did not observe any changes in resistance to the antibiotics used to treat the patient. The initial genetic diversity in the gut (2/12 isolates with SNPs) was higher than the diversity found in the lung (0/24 isolates with SNPs), and this difference was marginally significant (Z=2.05, P=.04). Two isolates from day 7 differed from the dominant ancestral strain by 5 SNPs and 3 indels. The high number of mutations suggests that these isolates were from a secondary intestinal colonization event by a closely related clone of ST17. This argument is further supported by the observation that these isolates lack the *ampD*H98Y mutation that was found in the ancestral strain.

### Fitness

Antibiotic resistance mutations are usually associated with fitness costs that can generate selection against resistance following antibiotic treatment [28, 29]. To better understand the link between antibiotic resistance and fitness in this infection, we measured the growth rate of all the lung isolates in nutrient-rich culture medium lacking antibiotics (Figure 4). We did not find any evidence that isolates with *oprD* or *wbpM* mutations had reduced fitness relative to the ancestral strain. Although this result is at odds with the established link between resistance and fitness, recent work has shown that *oprD* mutations do not generate a fitness cost, both *in vitro* and *in vivo* [30]. Given that o*prD* mutants were replaced by strains with mutations in *wbpM* or MexAB-OprM, we hypothesized that isolates with mutations in these genes would have higher fitness that *oprD* mutants. Consistent with our hypothesis, we found that isolates with mutations in *wbpM* (t19=2.02 one-tailed t-P=.0253) or MexAB-OprM (t19=2.34, one-tailed P=.0163) had higher fitness than *oprD* mutants. The low fitness of OprD mutants provides a simple explanation for the loss of high-level meropenem resistance following treatment, as meropenem sensitive MexAB-OprM mutants were able to replace competitively inferior OprD mutations.

**Figure 4.**
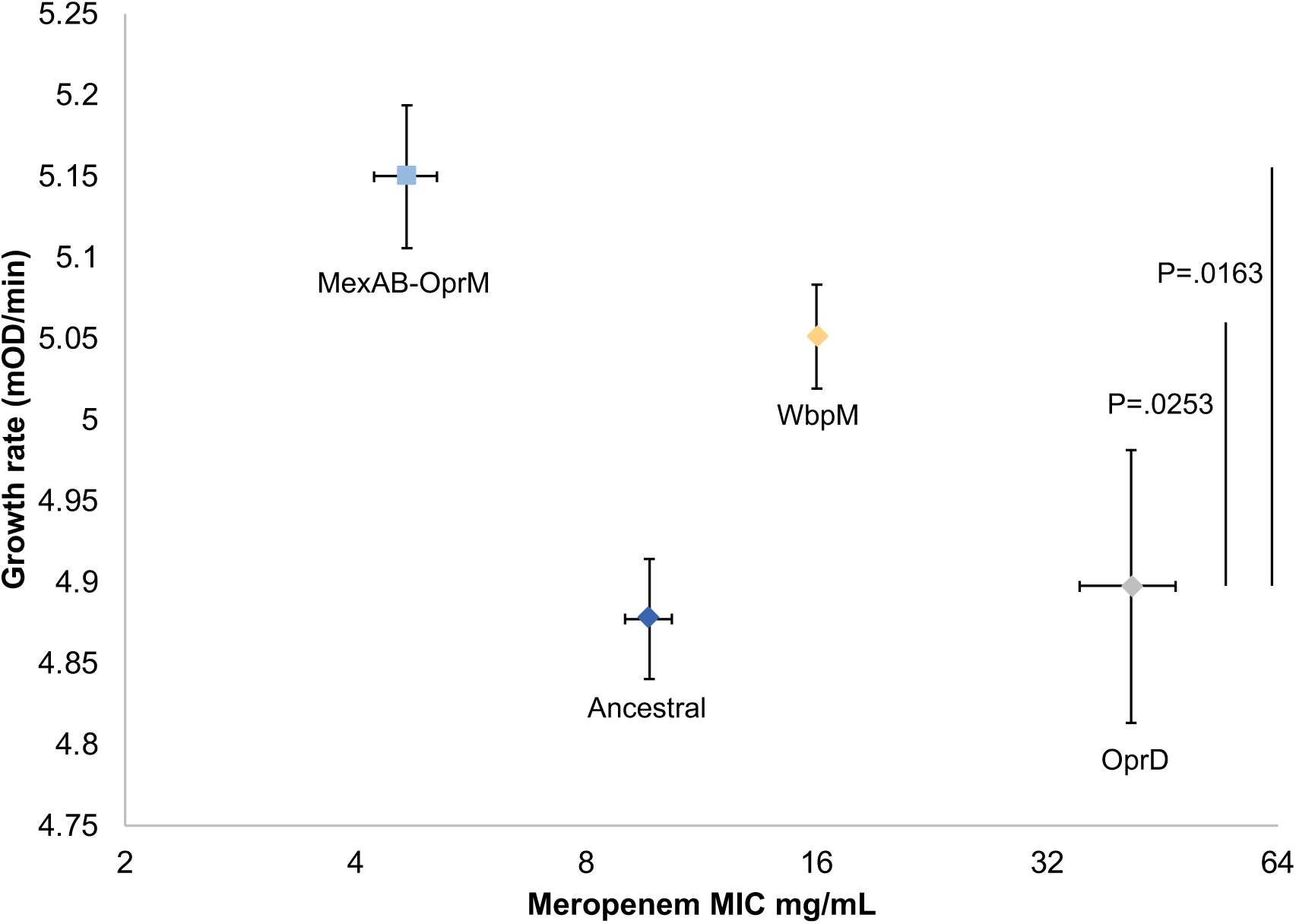
Fitness (growth rate) and meropenem resistance. The fitness and meropenem resistance of respiratory isolates according to mutated target (+/- s.e.m; n>5). Fitness was measured as log phase growth rate in culture medium lacking antibiotics. Elevated meropenem resistance was not associated with a fitness cost relative to ancestral isolates. However, both WbpM and MexAB-OprM mutants had high fitness relative to OprD mutants, as determined by one-tailed tests.

### Collateral responses to antibiotic treatment

There is growing interest in understanding the collateral changes in antibiotic resistance that emerge as a consequence of evolutionary changes to antibiotic treatment. For example, collateral sensitivity, which occurs when evolving increased resistance to one antibiotic confers increased sensitivity to alternative antibiotics, can be exploited to design combination therapies that restrict the evolution of resistance [31-33]. To investigate the collateral responses to meropenem treatment, we measured the resistance of our isolates to four alternative antibiotics with different mechanisms of action and resistance: ciprofloxacin (fluoroquinolone), gentamicin (aminoglycoside), ceftazidime and aztreonam (ß-lactams). Consistent with our previous data, we found no collateral changes in antibiotic resistance in gut isolates. In contrast, we found evidence of widespread, but subtle, variation in resistance to these alternative antibiotics across lung isolates.

Principal component analysis revealed 2 statistically independent axes of variation in this data set that reflect resistance to ciprofloxacin, gentamicin and ceftazidime (PC1; 47% of variation) and aztreonam (PC2; 25% of variation) (Figure 5). Isolates with mutations in the MexAB-OprM pump had reduced PC1 scores (Figure 5B; P<.0001, Dunnett’s test) and decreased colistin tolerance (Figure 5C; P=.0091, Dunnett’s test), reflecting the broad substrate range of this important pump.

**Figure 5.**
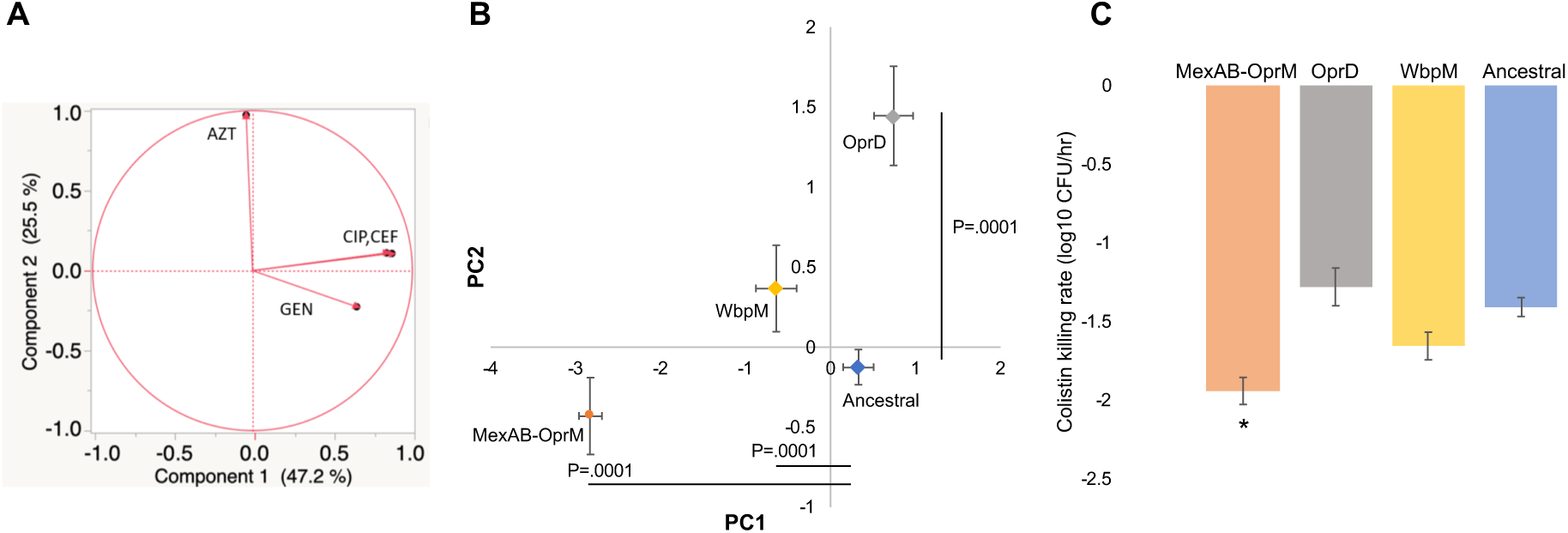
Collateral responses to antibiotic treatment. Panel A shows the results of principal component analysis of resistance to a panel of antibiotics that were not used to treat the patient, component 1 (PC1) and component 2 (PC2). Panel B shows the mean (+/- s.e.m.) principal component score for isolates according to different target mutations, and p values show the significant differences to the ancestral strain, as determined by Dunnett’s test. Panel C shows the colistin tolerance, as measured by the rate of cell killing at 2mg/L colistin. Altered colistin tolerance was only found in MexAB-OprM mutants, as determined by Dunnett’s test (P=.0091).

We found no evidence of collateral sensitivity associated with *oprD* mutations; rather, we found that *oprD* mutations conferred cross-resistance to aztreonam (i.e. PC2; Dunnett’s test P<.0001). Mutations in *wbpM*, on the other hand, generated sensitivity to a range of other antibiotics (PC1; P<.0001, Dunnett’s test) without creating cross-resistance to Aztreonam (PC2; P=.205, Dunnett’s test).

## Conclusion

### Summary

Nosocomial *P. aeruginosa* infections are often caused by strains that have high levels of antibiotic resistance, and carbapenem antibiotics and colistin have become key to the treatment of these infections [34-36]. Levels of resistance to colistin in *P. aeruginosa* remain low, but carbapenem resistance has increased to the point where the World Health Organisation has designated carbapenem resistant *P. aeruginosa* as a ‘critical priority’ for the development of new antibiotics. Combining clinical data (Figure 1), resistance phenotyping (Figure 2), genomics (Figure 3) and fitness assays (Figure 4) enabled us to produce a very high-resolution understanding of bacterial responses to meropenem therapy in a patient with a *P. aeruginosa* pneumonia, as summarized in Figure 6.

**Figure 6.**
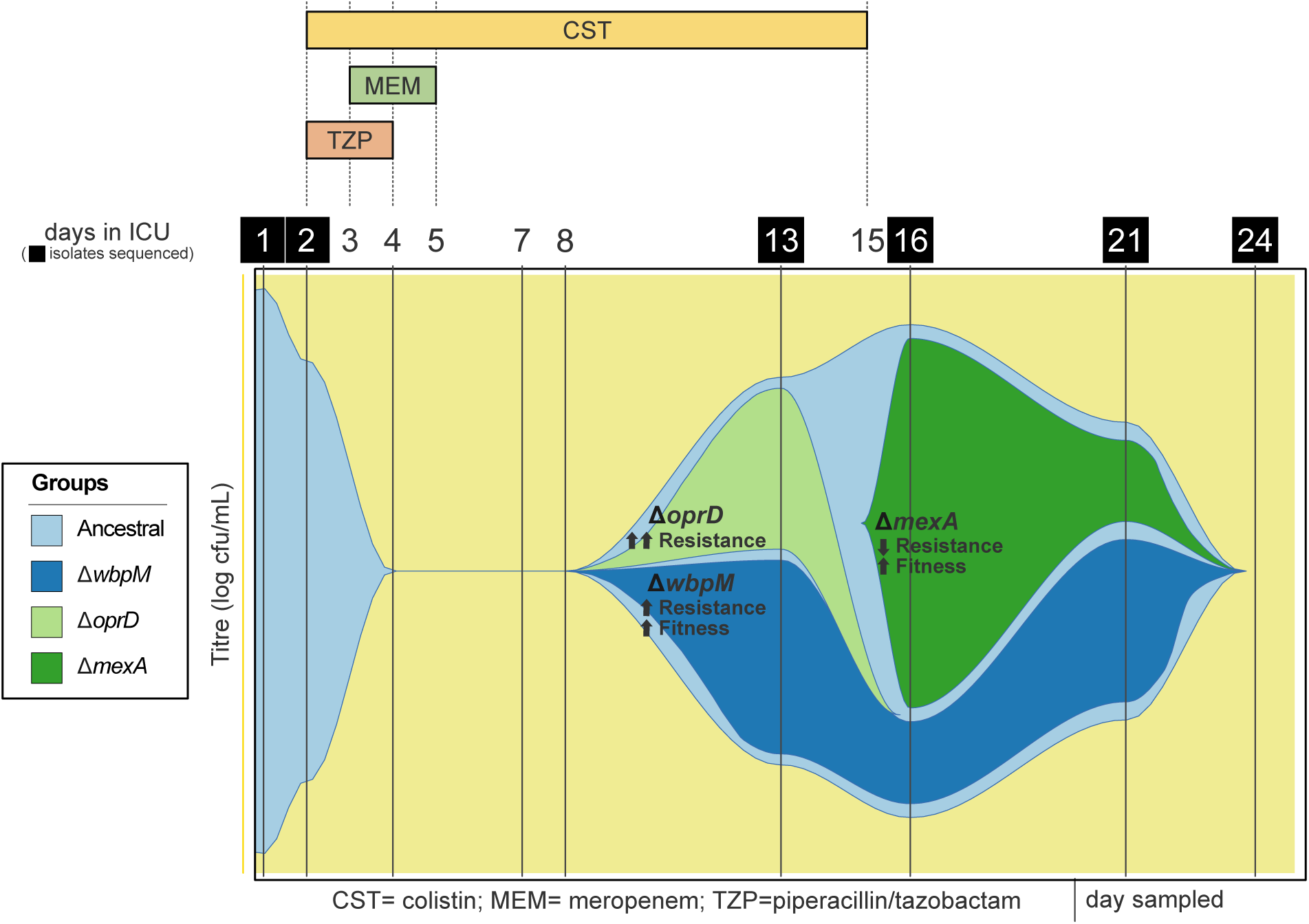
Graphical representation (Muller plot) of changes in the absolute abundance of the three main genotypes that we detected in lung samples. Annotations show changes in meropenem resistance and fitness relative to the ancestral strain.

### Responses to antibiotic treatment

We identified 3 independent evolutionary origins of meropenem resistance by mutations in *oprD* among a collection of only 107 isolates, including 2 singletons, implying that the true number of origins of OprD-mediated meropenem resistance in this patient was likely to have been much greater. We did not find any fitness costs associated with OprD resistance (see also [30]) or collateral sensitivity associated with *oprD* mutations, highlighting the incredible challenge of stopping *P. aeruginosa* from evolving high levels of carbapenem resistance via this simple mechanism.

In addition to this well-established mechanism of carbapenem resistance, we found evidence of selection for LPS biosynthesis (i.e. *wbpM*) mutations in response to meropenem treatment. Although the increase in resistance provided by *wbpM* mutations was modest (2x), *wbpM* mutations also increased fitness, allowing them to stably persist after meropenem treatment. Mutations that confer small increases in resistance are often regarded as being unimportant in clinical settings but in this case, the subtle increase in resistance provided by WbpM was sufficient to push the meropenem resistance of the ST17 clone that initiated the infection above the clinical breakpoint MIC. We suggest that further attention should be given to the role of these small effect mutations in antibiotic therapy, particularly for genes that can increase both resistance and fitness.

Colistin currently provides a last line of defence for the treatment of infections caused by Gram negative pathogens, and the rise of carbapenem resistance has resulted in increased use of colistin to treat *P. aeruginosa* infections [37]. The rapid decline in titre under colistin treatment observed between day 1 and 2 provides good evidence that colistin therapy was initially effective. However, *P. aeruginosa* recovered under colistin treatment without any associated increase in MIC or colistin tolerance. One possible explanation for these results is that the effective concentration of colistin in the lung decreased over time due to changes in drug pharmacokinetics. Alternatively, it is possible that adaptive changes in gene expression occurred in the lung increased phenotypic resistance to colistin in a way that cannot be captured by *in vitro* susceptibility assays, such as altered expression of the *arnBCADTEF* operon driven by two-component regulators such as PhoP/PhoQ, ColR/ColS or ParR/ParS [38]. This hypothesis could explain both why treatment was initially successful (i.e. rapid decline in *P. aeruginosa* titre under colistin treatment) and why colistin therapy fails to prevent the recovery of *P. aeruginosa* titre. The failure of colistin treatment in this patient is alarming, but it is not without precedence-colistin treatment of *P. aeruginosa* infections has a high failure rate, in spite of the fact that colistin resistant *P. aeruginosa* are very rare [39-41]. Our work highlights the need to better understand how *Pseudomonas* populations respond to colistin treatment during infections.

### Linking respiratory infection and intestinal carriage

Gut colonization with *P. aeruginosa* is associated with an increased risk of developing invasive lung infections [17] but the link between intestinal carriage of *P. aeruginosa* and lung infection remains poorly understood. Interestingly, genetic diversity was higher in the gut than in the lung at the onset of infection. One possible explanation for this observation is that both the lung and gut were independently colonized from a ‘source’ population of ST17 in the hospital environment. According to this explanation, the high diversity of gut isolates could have been caused by either early colonization followed by *in situ* diversification or by recurrent colonization events. In support of the latter scenario, we detected 2 gut isolates that reflect secondary colonization by a closely related lineage of ST17. Alternatively, it is possible that initial diversity in the lung was low because of a bottleneck associated with gut to lung transmission of *P. aeruginosa*. It is not possible to distinguish between these two hypotheses given our data, but our results highlight the need to better understand the population biology of intestinal and pulmonary populations of *P. aeruginosa*. A further challenge will be to understand the differential effects of antibiotics in the gut and lung. In contrast to the rapid evolution observed in the lung, we did not find any genetic or phenotypic evidence for selection in response to antibiotic treatment in the gut. The simplest explanation for the lack of response to treatment is that the intestinal population of *Pseudomonas* was not exposed to inhibitory concentrations of antibiotics, but it is also conceivable that the intestinal population did not evolve in response to treatment because of either a lack of available resistance mutations due to low population size or weak selection for resistance mutations due to elevated costs of resistance in the gut.

### Fitness costs and the fall of antibiotic resistance

It is well documented that antibiotic resistance tends to decline following treatment, but the underlying causes of this decline remain poorly understood [42]. Evolving antibiotic resistance tends to generate a fitness cost, suggesting that resistant strains should be replaced by superior competitors following antibiotic treatment [29]. However, experimental evolution studies show that resistance can be rapidly stabilized following antibiotic treatment by the spread of compensatory mutations that offset the cost of resistance [43, 44]. Although OprD-mediated resistance to meropenem did not generate a fitness cost *per se*, OprD mutants were rapidly replaced by high fitness strains with inactivating mutations in MexAB-OprM, a costly multi-drug efflux pump [30, 45]. Whereas compensatory evolution allows resistance to be stably maintained following antibiotic treatment, selection for ‘anti-resistance’ mutations resulted in reduced resistance to both meropenem and a spectrum of alternative antibiotics that were not used to treat this patient. Intriguingly, MexAB-OprM mutations are commonly detected in *P. aeruginosa* from patients with cystic fibrosis [46], suggesting that selection against this resistance may be a common feature of *Pseudomonas* infections. On the one hand, it is possible that antibiotic treatment creates conditions that generate selection for MexAB-OprM mutants. For example, by suppressing pathogen population density, antibiotic treatment may create conditions that generate strong selection for mutants with increased growth rate after antibiotic treatment ends. Alternatively, it is possible that MexAB-OprM mutations facilitate adaptation to the human lung in an opportunistic pathogen irrespective of antibiotic treatment.

### Conclusion and outlook

Although it has long been known that antibiotic treatment can drive the spread of resistance during infections, the underlying dynamics of this process remain poorly characterized, especially during acute infections by opportunistic and commensal pathogens. Our study was able to capture the evolutionary responses of a pathogen population to antibiotic treatment by characterizing the genetic and phenotype diversity present in the longitudinal samples taken from a single patient. The key insight from this is that natural selection can drive the incredibly rapid rise, and fall, of resistance. Previous work that has characterized the evolutionary dynamics of resistance in patients has relied largely on long-term sampling of chronic infections. Although our study has focused on a single patient, our findings suggest that infrequent sampling of pathogen populations may underestimate the rate of evolution of resistance because of the high rate at which resistant lineages rise and fall following treatment. Hopefully, future studies using our high-resolution approach across multiple patients will help to resolve this.

## Methods

### Clinical data

The patient was recruited as part of an observational, prospective, multicentre European epidemiological cohort study, ASPIRE-ICU (The Advanced understanding of *Staphylococcus aureus* and *Pseudomonas aeruginosa* Infections in Europe– Intensive Care Units, NCT02413242 ClinicalTrials.gov) that has been previously described [22]. This study enrolled subjects who were mechanically ventilated at ICU admission and with an expected length of hospital stay ≥48 hours. An assessment of four clinical criteria to establish a clinical diagnosis of ICU pneumonia (e.g. new blood culture drawn, new antibiotic use, new radiologic evidence, reason to suspect pneumonia) was performed daily; in case of at least one positive parameter, a combination of objective major and minor criteria was assessed to categorize subjects as having protocol pneumonia or not [22]. Data on antibiotic use in the two weeks preceding ICU admission and during the ICU stay were reported. During ICU stay, study samples (e.g. lower respiratory tract samples and peri-anal swabs) were obtained three times weekly in the first week, two times weekly in the three following weeks and on the day of diagnosis of protocol pneumonia and seven days after it.

### Sample collection and isolation

The respiratory samples and peri-anal swabs used in this study were collected within the ASPIRE-ICU study and are from a single patient at a Spanish hospital [22]. Respiratory samples were collected on the following visit days: 1 (the day of informed consent, 72h after ICU admission), 4, 7, and twice weekly for 30 days or until ICU discharge. In this case: day 10, 13, 16, 21, 23, 27. From patients who were diagnosed with pneumonia, additional respiratory samples were collected at the day of diagnosis and 7 days post-infection: day 2 and 8. Peri-anal swabs in skimmed milk medium and untreated respiratory samples were stored at -80 °C until shipment to the Central lab at the University of Antwerp and until further analysis. Semi-quantitative culture of peri-anal swabs was performed by inoculating the swabs directly on CHROMID *P. aeruginosa* Agar (BioMérieux, France) and blood agar (BBL®Columbia II Agar Base (BD Diagnostics, USA) supplemented with 5% defibrinated horse blood (TCS Bioscience, UK)). After incubation of 24 h at 37 °C, the growth of *P. aeruginosa* was evaluated in four quadrants. Plates without growth were further incubated for 48 h and 72 h.

Patient ETA samples were blended (30,000 rpm, probe size 8 mm, steps of 10 s, max 60 s in total), diluted 1:1 v/v with Lysomucil (10% Acetylcysteine solution) (Zambon S.A, Belgium) and incubated for 30 minutes at 37 °C with 10 s vortexing every 15 minutes. Thereafter, quantitative culture was performed by inoculating 10-fold dilutions on CHROMID *P. aeruginosa* Agar and blood agar using spiral plater EddyJet (IUL, Spain). Plates were incubated at 37 °C for 24 h and CFU/mL was calculated. Plates without growth were further incubated for 48 h and 72 h. Matrix-Assisted Laser Desorption Ionization-Time of Flight Mass Spectrometry (MALDI-TOF MS) was used to identify 12 colonies per sample which were stored at -80 °C until further use.

### Resistance phenotyping

All isolates were grown from glycerol stocks on Luria-Bertani (LB) Miller Agar plates overnight at 37°C. Single colonies were then inoculated into LB Miller broth for 18-20h overnight growth at 37°C with shaking at 225RPM. Overnight suspensions were serial diluted by 1:10000 fold to ∼ 5 x 10^5^ CFU/mL. Resistance phenotyping was carried out as minimum inhibitory concentration (MIC) testing via broth microdilution as defined by EUCAST recommendations [47, 48], with the alteration of LB Miller broth for growth media and the use of *P. aeruginosa* PAO1 as a reference strain [49]. Three biological replicates of each isolate were used to calculate MIC, and the MIC of each isolate was calculated from the median.

### Colistin tolerance assay

All isolates were grown from glycerol stocks on LB Miller Agar plates overnight at 37°C. Each culture was grown from a single, randomly selected colony, inoculated into 200ul of LB Miller and grown over 18-20h at 37°C with shaking at 225RMP. Overnight cultures were diluted in phosphate saline buffer to a final concentration of approximately 1×10^6^ CFU/mL, further verified by total viable count, and grown in 200ul LB Miller with or without the addition of 2 mg/L colistin. To avoid colistin carry-over, cultures were diluted at least 10-fold and plated on LB Miller agar after 0, 1, 2, 4 or 8 hours. This assay was carried out in using a randomized block experimental design, and we analysed 5 replicates of 11 randomly selected isolates and 5 replicates of a PA01 control in each block. The order and position of each isolate on the experimental plates was selected through randomisation, using ‘sample’ command without replacement in R [50]. We used linear regression of mean log viable cell titre against time to calculate a death rate for each isolate (typically this involved data from 0-2 hours of incubation). In no case did we observe bi-phasic killing kinetics.

### Characterization of the *wbpM* mutant

In order to determine the effect of the *wbpM* in resistance, meropenem MICs were determined by EUCAST broth microdilution in triplicate experiments for wild-type reference strain PA14 and its *wbpM* isogenic knock out derivative obtained from an available transposon mutant library [51].

### Gene expression

The expression of the genes encoding the chromosomal β-lactamase AmpC (*ampC*) and MexAB-OprM efflux pump (*mexB*) was determined from late-log-phase LB broth cultures at 37°C and 180 rpm by real-time RT–PCR with an Bio-Rad (CFX Connect Real-Time System), as previously described [52].

### Long-read sequencing

Four isolates were sequenced with the Pacific Biosciences platform using single molecule chemistry on a SMRT DNA sequencing system. Coverage ranged from 122X - 171X. Resulting sequencing reads were assembled using canu v. 1,8 indicating a genome size of 7Mb and using raw error rate of 0.300, corrected error rate of 0.045, minimum read length of 1,000 bases, and minimum overlap length of 500 [53]. Canu assemblies were circularized using circlator v.1.5.5 testing kmer sizes 77, 87, 97, 107,117, and 127, minimum merge length of 4,000, minimum merge identity of 0.95, and minimum contig length of 2,000 [54].

### Illumina Sequencing

All isolates were sequenced in the MiSeq or NextSeq illumina platforms yielding a sequencing coverage of 69X – 134X. Raw reads were quality controlled with the ILLUMINACLIP (2:30:10) and SLIDINGWINDOW (4:15) in trimmomatic v. 0.39 [55]. Quality controlled reads were assembled for each isolate with SPAdes v. 3.13.1 with default parameters [56]. These assemblies were further polished using pilon v. 1.23 with minimum number of flank bases of 10, gap margin of 100,000, and kmer size of 47 [57]. Resulting contigs were annotated based on the *P. aeruginosa* strain UCBPP-PA14 [58] in prokka v. 1.14.0 [59].

### Variant calling

To identify pre-existing resistance mutations that were present at the start of the infection reads for each of the isolates were mapped to the *P. aeruginosa* PAO1 reference genome (GenBank accession: NC_002516.2) with Bowtie 2 v2.2.4 [60] and pileup and raw files were obtained by using SAMtools v0.1.16 [61] and PicardTools v1.140 [62], using the Genome Analysis Toolkit (GATK) v3.4.46 for realignment around InDels [63]. From the obtained raw files, SNPs were extracted if they met the following criteria: a quality score (Phred-scaled probability of the samples reads being a homozygous reference) of at least 50, a root-mean-square (RMS) mapping quality of at least 25 and a coverage depth of at least 3 reads; excluding all ambiguous variants. As well, MicroInDels were extracted from the total pileup files when meeting the following: a quality score of at least 500, a RMS mapping quality of at least 25 and support from at least one-fifth of the covering reads. Filtered files were eventually annotated with SnpEff v4.2 and SNPs and InDels located in a set of genes known to be involved in *P. aeruginosa* chromosomal antibiotic resistance were extracted [64-66].

To identify mutations and gene gain/loss during the infection, short-length sequencing reads from each isolate were mapped to each of the four long-read de novo assemblies using bwa v. 0.7.17 using the BWA-MEM algorithm [67]. Preliminary SNPs were identified with SAMtools and BCFtools v. 1.9 [61]. Low-quality SNPs were filtered out using a two-step SNP calling pipeline which first identified potential SNPs using the following criteria: 1. Variant phred quality score of 30 or higher, 2. At least 150 bases away from contig edge or indel, and 3. 20 or more sequencing reads covering the potential SNP position [7]. In the second step, each preliminary SNP was reviewed for evidence of support for the reference or the variant base; at least 80% of reads of phred quality score of 25 or higher were required to support the final call. An ambiguous call was defined as one with not enough support for the reference or the variant, and, in total, only one non-phylogenetically informative SNP position had ambiguous calls. Indels were identified by the overlap between the HaplotypeCaller of GATK v. 4.1.3.0 [68] and breseq v. 0.34.0 [69]. The variable genome was surveyed using GenAPI v. 1.0 [70] based on the prokka annotation of the short-read de novo assemblies. The presence or absence of genes in the potential variable genome was reviewed by mapping the sequencing reads to the respective genes with BWA v.0.7.17 [64-66].

### Growth rate assays

All isolates were grown from glycerol stocks on LB Miller Agar plates overnight at 37°C. Single colonies were then inoculated into LB Miller broth for 18-20h overnight growth at 37°C with shaking at 225RPM. Overnight suspensions were serially diluted to an OD595 of ∼ 0.05 and placed within the inner wells of a 96-well plate equipped with a lid. To assess growth rate, isolates were then grown in LB Miller broth at 37°C and optical density (OD595nm) measurements were taken at 10-minute intervals in a BioTek Synergy 2 microplate reader set to moderate continuous shaking. Growth rate was calculated as the maximum slope of OD versus time over an interval of 10 consecutive readings, and we visually inspected plots to confirm that this captured log-phase growth rate. For lung isolates, growth rate was calculated from the mean of 10 biological replicates, and for peri-anal isolates, growth rate was calculated as the mean from > 4 biological replicates

## Acknowledgements

This research project was by Wellcome Trust Grant (106918/Z/15/Z) through the Innovative Medicines Initiative Joint Undertaking under COMBACTE-MAGNET (Combatting Bacterial Resistance in Europe - Molecules against Gram-negative Infections, grant agreement n° 115737) and COMBACTE-NET (Combatting Bacterial Resistance in Europe - Networks, grant agreement n° 115523), resources of which are composed of financial contribution from the European Union’s Seventh Framework Programme (FP7/2007-2013) and EFPIA companies’ in kind contribution. We thank the Oxford Genomics Center (funded by Wellcome Trust Grant 203141/Z/16/Z) for the generation and initial processing of Illumina sequence data.

## Notes

### Competing Interest Statement

The authors have declared no competing interest.

